# The Evolution of Phenotypic Plasticity when Environments Fluctuate in Time and Space

**DOI:** 10.1101/395137

**Authors:** Jessica G. King, Jarrod D. Hadfield

## Abstract

Most studies have explored the evolution of plasticity when the environment, and therefore the optimal trait, varies in time or space. When the environment varies in time and space we show that genetic adaptation to temporal fluctuations depends on the between-generation autocorrelation in the environment in exactly the same way that genetic adaptation to spatial fluctuations depends on the probability of philopatry. This is because both measure the correlation in parent-offspring environments and therefore the effectiveness of a genetic response to selection. If the capacity to genetically respond to selection is stronger in one dimension (e.g. space) then plasticity mainly evolves in response to fluctuations in the other dimension (e.g. time). If the relationship between the environments of development and selection are the same in time and space then the evolved plastic response to temporal fluctuations is useful in a spatial context and genetic differentiation in space is reduced. However, if the relationship between the environments of development and selection are different then the optimal level of plasticity is different in the two dimensions. In this case the plastic response that evolves to cope with temporal fluctuations may actually be maladaptive in space. This can result in the evolution of hyperplasticity or negative plasticity, the effects of which are mitigated by spatial genetic differentiation. However, genetic differentiation acts in opposition to plasticity resulting in counter-gradient variation. These results highlight the difficulty of making space-for-time substitutions in empirical work but identify the key parameters which need to be measured in order to test whether space-for-time substitutions are likely to be valid.

## Introduction

Phenotypic plasticity is the ability of a single genotype to produce different phenotypes when exposed to different environmental settings. It is a ubiquitous feature of organisms (West-Eberhard, 2003; Pigliucci, 2001) that can be broadly divided into adaptive and non-adaptive categories (Ghalambor et al., 2007). Adaptive plasticity arises as an evolved ‘active’ response to environmental fluctuations that allows organisms to produce phenotypes better matched to their environment. It remains the focus of much empirical work and studies continue to be published that give new insights and provide pivotal tests of key ideas (Dey et al., 2016; Huang and Agrawal, 2016; van Buskirk, 2017). This empirical work is supported by a large body of theoretical work that has either considered scenarios where the environment fluctuates in time in a single population, or in space across multiple populations connected by migration. In the absence of intrinsic costs to plasticity, and when the environment of development (the environmental variable that induces the plastic response) is a perfect predictor of the environment of selection (the environmental variable that determines the optimal trait value), perfect plasticity is known to evolve (Via and Lande, 1985). In this scenario, plasticity allows organisms to perfectly adjust to environmental conditions and any genetic differentiation in time or space is lost.

When the environments of development and selection are not perfectly correlated (Moran, 1992), the evolved plastic response is shallower than the perfect response and equal to the regression of the optimum on the environment of development (Gavrilets and Scheiner, 1993; de Jong, 1999; Tufto, 2000). In what follows we will refer to this regression as the DO-regression, the magnitude of which can be interpreted as cue reliability. When there is some cue unreliability, genetic differentiation can occur because a shortfall exists between the optimal trait value and the trait value induced by the plastic response to the environment (de Jong, 1999; Tufto, 2000). The effects of this shortfall can be mitigated when the optima of parents are similar to those of their offspring because trait values can partly track fluctuations in the optimum through genetic adaptation. In what follows we will refer to the similarity of parent and offspring optima as the PO-regression. The PO-regression will be high if the environment changes little between generations and changes little over the spatial scale at which individuals disperse; the PO-regression will be high when temporal and spatial autocorrelation is high. Theoretical models have quantified the expected degree of genetic tracking in temporally (Michel et al., 2014) and spatially (Hadfield, 2016) autocorrelated environments given some predefined level of plasticity. Intuitively, the amount of genetic tracking is found to increase when there is substantial genetic variance and weak plasticity, particularly when selection is strong around a widely fluctuating and highly autocorrelated optimum (as in models without plasticity; Slatkin and Lande, 1976; Lande and Shannon, 1996). The degree of genetic tracking can be measured as the covariance between genetic values and the optimum (Blanquart et al., 2012), a quantity shown to have the same form in continuous time (Michel et al., 2014) and continuous space (Hadfield, 2016) models when autocorrelation is scaled to generation-time or dispersal distance, respectively (Hadfield, 2016). The analytical models of Michel et al. (2014) and Hadfield (2016) treated plasticity as a fixed rather than evolving parameter. However, Tufto (2015) showed that in a discrete time model, where plasticity is free to evolve, the same results hold and here we confirm that this is also true for discrete space. Although these results clearly show that the evolved level of plasticity determines the degree to which genetic tracking occurs, it remains less clear to what extent the capacity to genetically track environmental change determines the level of plasticity (Tufto, 2015).

In many previous models it has been hard to address this question because the environments of selection and development are often treated as a single environmental variable but experienced at different life-stages. This causes the PO- and DO-regressions to depend on the same parameters, making it unclear whether it is the capacity to genetically track environmental change or cue reliability that is driving the evolution of plasticity. For example, with low migration, parents and offspring experience more similar environments (the PO-regression is large), but the environments of selection and development are also more similar because more individuals are subject to selection where they develop (the DO-regression is also large) (de Jong, 1999). de Jong (1999) developed a discrete spatial model where these effects could be separated, and concluded that the plastic slope did not depend on the capacity to genetically track environmental change but evolved to the DO-regression; an identical result to that in Gavrilets and Scheiner (1993) in which the PO-regression is implicitly zero and therefore no capacity to genetically track environmental change exists. In contrast, Tufto (2000) developed a similar spatial model and found (Equation 15) that the slope was shallower than that in Gavrilets and Scheiner (1993). Although no interpretation of this result was given, the degree to which the plastic slope became shallower scaled positively with parameters that increase the rate of genetic tracking, suggesting that the capacity to adapt may impact on the evolution of plasticity. Subsequent work suggests that this result arises because in Tufto’s (2000) model the environment of development varies over individuals within populations which elevates the phenotypic variance in the trait when plasticity exists (Tufto, 2015). Under stabilising selection this increase in phenotypic variance imposes a cost on plasticity and so it evolves to be shallower when genetic tracking is possible (Tufto, 2015). This effect was implicitly ignored in de Jong (1999) due to a weak selection approximation, and in general we may expect the effect to be small. However, a direct cost to plasticity may be stronger (van Tienderen, 1991; DeWitt et al., 1998) and models that include such costs result in reduced plastic responses (van Tienderen, 1997; Lande, 2014). Intuition suggests that the degree to which the plastic response is reduced in the presence of costs should depend on the capacity of the population to genetically track environmental change and here we confirm that this is the case.

The effects of costs and cue unreliability are now well understood and provide a compelling explanation for why adaptive plastic responses are generally shallower than the perfect response. However, they fail to explain situations where plastic responses are steeper or in the opposite direction to the perfect response, called hyperplasticity or negative plasticity, respectively. Although these phenomena may be erroneously identified if the plastic response is determined by multiple cues only one of which has been mea-sured (Chevin and Lande, 2015), reciprocal transplant or experimental evolution studies offer robust tests and many convincing examples of hyperplasticity exist (Conover et al., 2009; Huang and Agrawal, 2016). Explanations for hyperplasticity usually invoke nonadaptive plasticity (Levins, 1968; Conover and Schultz, 1995) or adaptive plasticity that has become maladaptive by a sudden change in the environment (Van Asch et al., 2013; Cenzer, 2017). Often, these putatively maladaptive plastic responses are accompanied by compensatory genetic changes that have evolved to bring phenotypes closer to their optima (Levins, 1968), resulting in counter-gradient variation (Conover and Schultz, 1995; Grether, 2005).

To date, most studies have considered either temporal or spatial variation in the environment. However, Scheiner (2013) implemented a simulation model incorporating both. This work challenged the idea that spatial and temporal heterogeneity are substitutable and showed that plasticity evolves more easily in the presence of spatial heterogeneity compared to temporal heterogeneity. This work also demonstrated that, rather than being maladapative, hyperplasticity could evolve under extreme patterns of temporal variation in the environment (see Scheiner and Holt, 2012, also) due to the evolution of bet-hedging. However, as in many previous models, the environments of development and selection were the same variable experienced at different life stages causing the DO-regression to be inextricably tied to dispersal and generation-time, and as a consequence the PO-regression. Here we separate these two phenomena making it easier to understand how temporal and spatial fluctuations interact. We show that environmental heterogeneity in time *is* substitutable for that in space when spatial and temporal parameters are expressed in terms of spatial and temporal PO/DO-regressions. The equation for the equilibrium mean plasticity is pleasingly symmetric with respect to temporal and spatial parameters, and asymmetries only appear when temporal and spatial PO/DO-regressions take different values. If different enough, they can generate adaptive hyperplasticity (or negative plasticity) even in the absence of bet-hedging.

## Methods

### Model foundations

We consider a population composed of an infinite number of islands of infinite size, all of which exchange migrants at the same rate *m*. Islands differ in two environmental variables, one of which is a cue responsible for the plastic development of a trait (environment of development) and the other determines the selective consequences of expressing a particular trait value (environment of selection). In addition to spatial variation, both environmental variables fluctuate stochastically over time within each island according to an autoregressive process.

The order of events in the population is 1) fertilisation, 2) development, 3) selection, 4) gametogenesis and 5) migration. Phenotypes are assessed after development but before selection. The phenotype of individual *j* from island *i* at time *t* is a linear function of the environment of development (*D*_*it*_),

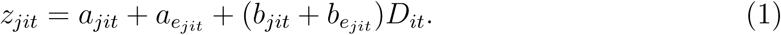

The intercept represents the component of the individual’s phenotype that is fixed across environments, with a genetic (*a*_*jit*_) and a non-genetic (*a*_*ejit*_) component. The slope determines how the phenotype responds to environmental variation, again with a genetic (*b*_*jit*_) and a non-genetic (*b*_*e* *jit*_) component. The variance in these components within an island at a specific point in time are assumed to be constant and are designated *G*^*aa*^, *E*^*aa*^, *G*^*bb*^ and *E*^*bb*^ respectively. We assume intercepts and slopes are uncorrelated, which is expected to evolve under stabilising selection (Lande, 2009), and that the environmental components have zero mean.

The optimal phenotype is assumed to depend linearly on the environment of selection (*S*_*it*_):

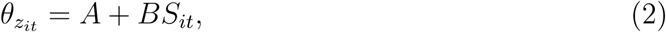

where intercept *A* represents the optimal phenotype in the reference (average) environment, and slope *B* the environmental sensitivity of the optimal phenotype (Chevin et al., 2010).

We assume that the environmental variables can be decomposed into separable spacetime processes of the form (for the environment of development)

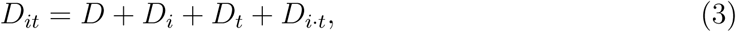

where *D* denotes the grand mean, *D*_*i*_ the deviation of island *i* from the grand mean (averaged over time), *D*_*t*_ the deviation at time *t* from the grand mean (averaged over islands) and *D*_*i·t*_ the deviation specific to a time and place. Time is measured in units of generations.

Spatial components of the environmental variables, *D*_*i*_ and *S*_*i*_, are assumed independent and identically distributed with corresponding variances 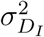 and 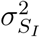, as are the space-time interaction components with variances 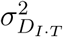 and 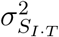 Temporal components are assumed to fluctuate according to an autoregressive process with stationary variances 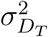 and 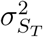 and a common autocorrelation parameter *α*_*T*_. The environments of selection and development are assumed to be linearly associated, with the regression of the environment of selection on the environment of development being *κ*_*I*_ in space, *κ*_*T*_ in time and *κ*_*I·T*_ at a specific place and time. The DO-regression is defined as *Bκ* and can be different in time and space if *κ*_*I*_ ≠ *κ*_*T*_.

The fitness of an individual on island *i* at time *t* is described by two independent Gaussian fitness functions. For the trait, the optimum of the fitness function is *θ*_*zit*_ and its width is *ω*_*z*_. For the slope, the optimum of the fitness function is 0 and its width is *ω*_*b*_ such that the absolute magnitude of plasticity is costly (van Tienderen, 1997; Lande, 2014; Kuijper and Hoyle, 2015) and can be thought of as a maintenance cost (DeWitt et al., 1998). Under this model, the strength of stabilising selection acting on the phenotype is 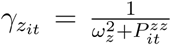 where 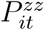 is the phenotypic variance on island *i* at time *t*. Likewise the strength of stabilising selection acting on the slope is 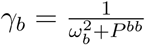

In what follows it will be useful to express migration by its opposite, the probability of philopatry; *α*_*I*_ = 1 - *m*. We choose this symbol due to its analogy with the temporal autocorrelation parameter *α*_*T*_, and note that the correlation between *D*_*i*_ (or *S*_*i*_) of parents and offspring is *α*_*I*_ in the same way that the correlation between *D*_*t*_ (or *S*_*t*_) of parents and offspring is *α*_*T*_. We refer to *α*_*I*_ and *α*_*T*_ collectively as PO-regressions, which can clearly be different in time and space. The correlation between *D*_*i·t*_ (or *S*_*i·t*_) of parents and offspring is zero (i.e. *α*_*I·T*_ = 0), because the deviations are unique to a specific generation and place, and so it is not possible to adapt to this source of variation. In the SI we discuss the likely consequences of allowing within-island temporal autocorrelation for *D*_*i·t*_ and *S*_*i·t*_ such that *α*_*I·T*_*≠* 0.

It should also be noted that the environments of development and selection have a common autocorrelation parameter *α*_*T*_ because we imposed it; it simplifies the analysis and makes the temporal model more comparable to the spatial model where *α*_*I*_ has to be common to both environmental variables because it is only a function of dispersal probability. A continuous space model, like the continuous time model, would allow *α*_*I*_ to be different for the two environmental variables because it would then depend on both dispersal distance and the spatial autocorrelation in the environment, which may differ between the environments of development and selection. In the SI we discuss the likely effect of assuming *D*_*t*_ and *S*_*t*_ have the same autocorrelation.

### Equilibrium Conditions

We are interested in obtaining equilibrium distributions for the mean intercept and slope on each island at each time (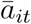 and 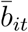). However, it is not possible to get analytical solutions for the model without making some additional assumptions and approximations. Throughout, we assume that *γ*_*zit*_ is constant in time and space (and therefore denoted as *γ*_*z*_). This latter approximation will hold if there is weak selection and/or if variation in the slopes is small. We also assume that variation in the mean slope over time within an island is small, which will be true if *G*^*bb*^ is small and/or temporal fluctuations are weak and not strongly autocorrelated. We relax these assumptions in a simulation model to assess the robustness of our conclusions.

As with the environmental variables we can write the mean reaction norm components as (for the intercept):

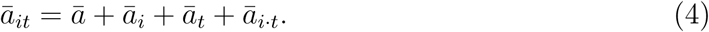

At equilibrium, we find the time-averaged mean intercept in island *i* is

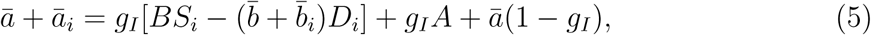

where 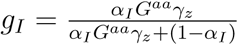 takes values between 0 and 1, where 0 indicates no capacity to genetically track spatial fluctuations in the optimum and 1 indicates complete capacity to genetically track the optimum. It increases when the probability of philopatry, the genetic variance in the intercept and/or the strength of stabilising selection around the optimum increase.

The time-averaged mean slope in island *i* is

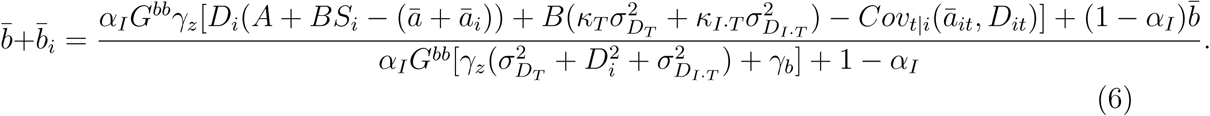

Equation 6 includes a term for the covariance between the mean intercept and the environment of development over time within island 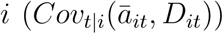. At equilibrium, this covariance is

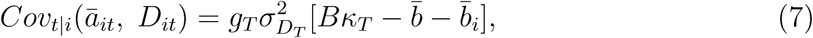

where 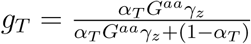 and has the same form as *g*_*I*_.

To obtain solutions for this system of equations, we also need expressions for 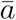 and 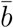, which are the expectations of Equations 5 and 6 over islands. In both cases, we can take a Taylor expansion around the mean environmental variables. For the mean intercept, a first order expansion is exact. For the mean slope, an exact expression is not obtainable and so we use a second order approximation (Tufto, 2000). The solutions to these equations are given in the results.

### Simulations

To test how accurate our approximations are, we simulated the process for 15,000 generations using a population of 1,000 islands. The first 5,000 generations were discarded to allow the process to reach equilibrium. A range of parameter values were used and are detailed in the results section and SI. The simulation was written in R and the code is available in the SI.

## Results

When solving Equation 5 with respect to the mean environment of development, the grand mean intercept is

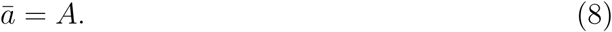

Substituting Equation 8 into 5, and solving for 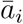 yields

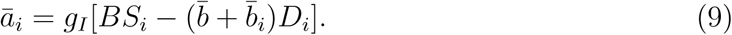

where the shortfall of the plasticity-induced phenotype from the optimum (the term in square brackets) is weighted by the capacity to genetically track changes in the optimum through space (*g*_*I*_).

Expressions for the mean slope and island slope deviations, and consequently island intercept deviations (Equation 9), are extremely complex and therefore only explored graphically. Before discussing them, it will be instructive to explore the solutions when *G*^*bb*^*→* 0. As this limit is approached, there is sufficient genetic variance in the slope for it to evolve to an equilibrium, but once at equilibrium spatial and temporal fluctuations are negligible. The mean slope is then

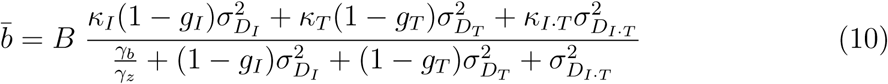

and spatial differentiation in the slope disappears, such that 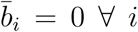. Equation 10 shows that the effects of spatial and temporal variation on the evolution of plasticity are symmetric, and that steeper slopes are favoured when the cost of plasticity is small, the capacity to genetically track environmental change is low and cue reliability is high. The role of variation specific to a time and place also has the same form, but because these fluctations are uncorrelated between generations *g*_*I·T*_ is effectively zero. At the extreme, when *κ*_*I*_ = *κ*_*T*_ = *κ*_*I·T*_ and there is no cost to plasticity (*γ*_*b*_ = 0), the mean slope is equal to *Bκ* (the DO-regression; Gavrilets and Scheiner, 1993), but becomes shallower as the cost increases and the capacity to genetically track environmental change becomes stronger. Under the same limit, *G*^*bb*^ *→* 0, the temporal covariance between the mean intercept and the environment of selection is given as

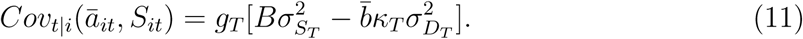

Tufto (2015) developed a model for the evolution of plasticity in a temporally autocorrelated environment, where the environments of development and selection are the same environmental variable separated by time *τ*. If we set spatial heterogeneity to 0 in our models, and note that 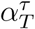 is equivalent to *κ*_*T*_ under this scenario, Equation 11 becomes equivalent to that in Tufto (2015, Equation 4c). The spatial covariance between the intercept and the environment of selection has the same form,

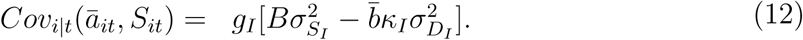

Equations 11 and 12 can be interpreted as measures of temporal and spatial (local) adaptation (Blanquart et al., 2012).

In the following graphical exploration of the solutions (where *G*^*bb*^ does not tend to zero) we focus primarily on spatial patterns rather than temporal patterns because they have been the focus of more empirical work. However, given the symmetrical effects of time and space the results can be directly applied to temporal patterns (Grether, 2005). In addition, we assume 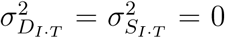 for ease of interpretation. If fluctuations specific to a time and place did exist, the plastic slope would be pulled toward *κ*_*I·T*_ because it is not possible to genetically track these fluctuations. Figure 1 illustrates the equilibrium solutions for the intercept, slope and phenotype as functions of the environment of selection across islands and the probability of philopatry. The left column portrays a model where only spatial variation exists. Panel a) shows genetic differentiation between populations increases as individuals become more philopatric; in the absence of migration, populations become perfectly adapted to local conditions. Panel b) shows that, when there is no philopatry, plasticity evolves to its maximum, which is slightly less than *κ*_*I*_ because of the cost to plasticity. For any given level of philopatry, there is slight genetic differentiation in the slope, which is shallowest in the average environment and steepest towards the extremes. This result has been shown using simulations (Scheiner, 1998) and analytically (Tufto, 2000), and arises because the genetic covariance between the slope and phenotype is higher in extreme environments and so the correlated response in the slope to selection on the phenotype is greater (Lande, 2009). The effects of plasticity and genetic differentiation combine to produce phenotypes that track the optimum closely, particularly when there is complete philopatry (Panel c)). The right column includes temporal fluctuations with the same properties as spatial fluctuations. Here, genetic differentiation between populations is reduced, even under complete philopatry (Panel d)), and plasticity plays a greater role in tracking spatial variation (Panel e)). This arises because plasticity that evolves to deal with temporal fluctuations is capable of also tracking some of the spatial variation.

**Figure 1:**
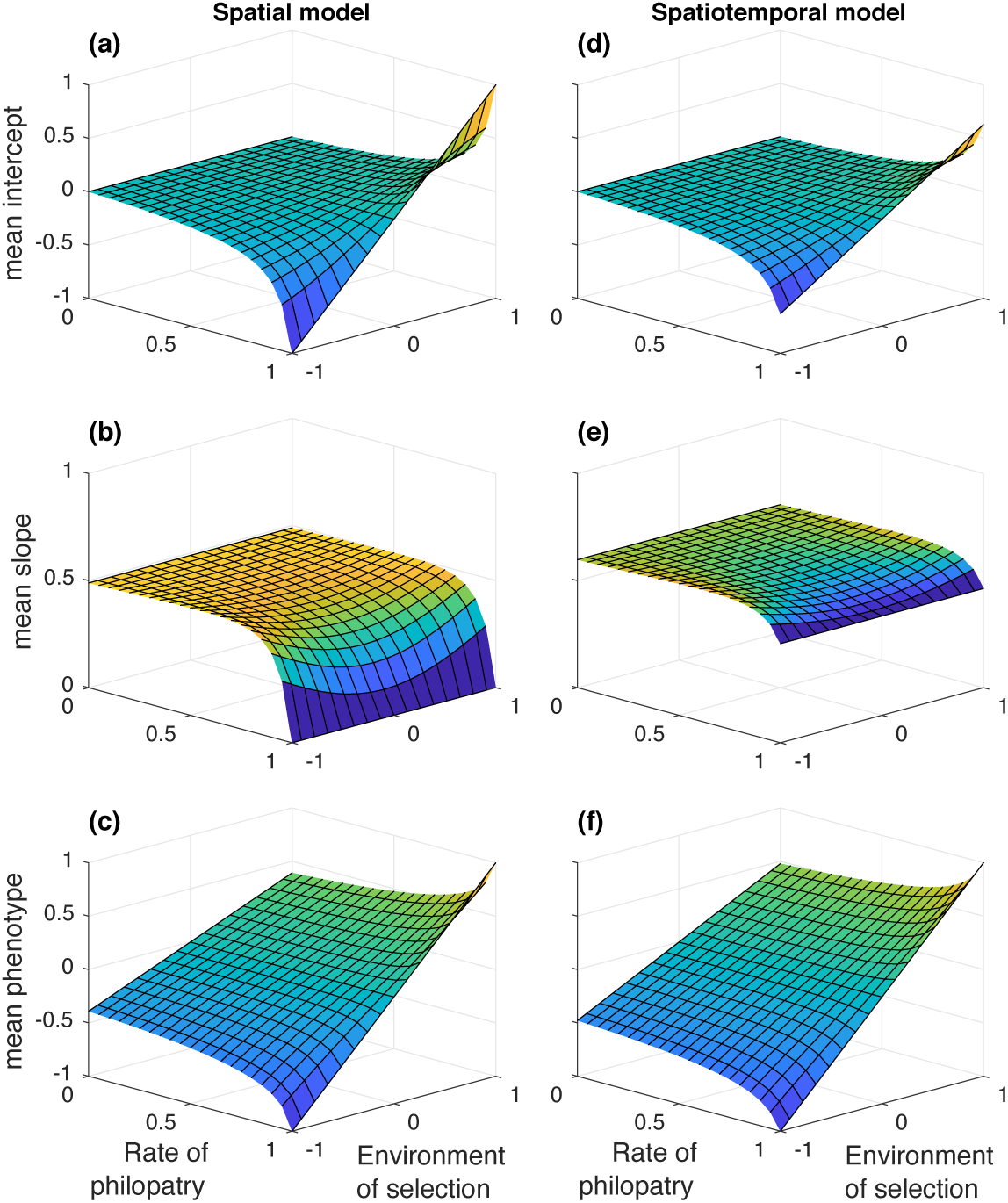
Island mean reaction norm components averaged over time and the environment of development, as functions of the spatial component of the environment of selection *S*_*i*_ and the probability of philopatry *α*_*I*_, with the intercepts being represented in the first row (a,d), the plastic slopes in the second row (b,e) and the phenotype in third row (c,f). The left column (a-c) represents a model where only spatial variation exists, such that any environmental variation in time is absent 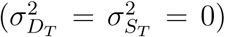, and the right column (d-f) a model where spatial and temporal variation exist simultaneously 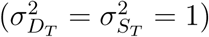. From the assumption that environmental fluctuations specific to a time and place are zero,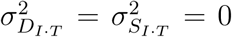. The remaining fixed parameters are 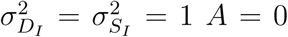, *B* = 1, *G*_*aa*_ = *G*_*bb*_ = *E*_*aa*_ = *E*_*bb*_ = 1, *ω*_*z*_ = 1, *ω*_*b*_ = 3, *κ*_*I*_ = *κ*_*T*_ = 0.8, *α*_*T*_ = 0.5.

*Figure 1 here*

Figure 2 illustrates how the mean slope (left column) and the spatial association between intercept and the environment of selection (right column) change as a function of various model parameters. Panels a) and b) show that the mean slope is affected in exactly the same way by temporal and spatial parameters. Panel c) illustrates a scenario where temporal parameters are fixed but the equivalent spatial parameters are allowed to vary. When there is no migration (*α*_*I*_ = 1), genetic tracking of spatial fluctuations is perfect such that selection on plasticity is determined solely by temporal parameters and does not depend on the spatial DO-regression. As migration increases, spatial genetic tracking becomes harder and plasticity evolves to also cope with spatial fluctuations. In the absence of a cost, the mean slope evolves to be intermediate between the spatial and temporal DO-regressions, and is pulled towards the spatial DO-regression as migration increases and spatial genetic tracking becomes harder.

**Figure 2:**
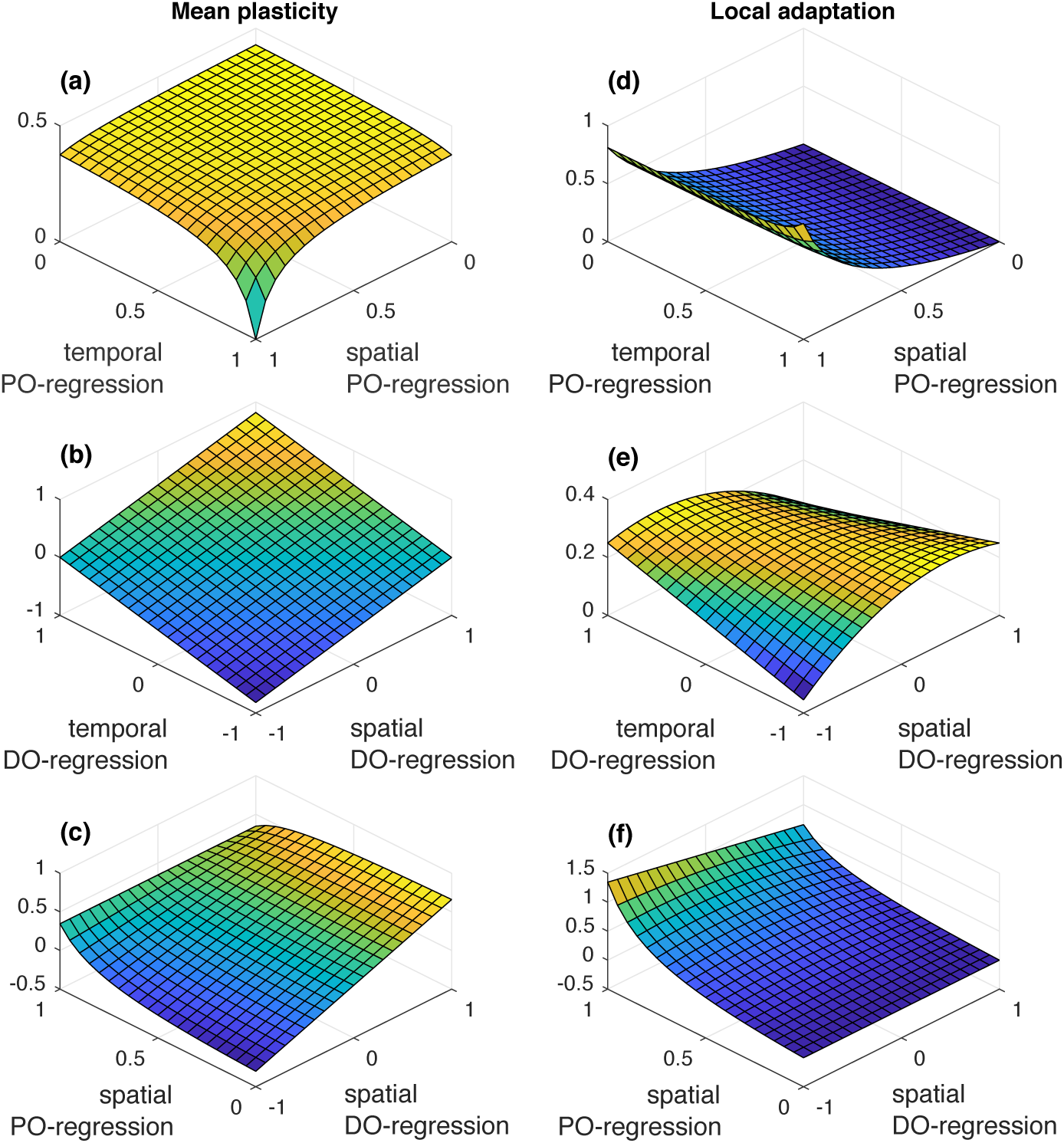
Mean plasticity (left column) and local adaptation (right column) as functions of model parameters. a) and b) show that the spatial and temporal parameters have a symmetric effect on mean plasticity and c) demonstrates what happens when the spatial parameters are allowed to vary but the temporal parameters are fixed (*κ*_*T*_ = 0.5, *α*_*T*_ = 0.5 and 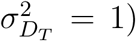. How genetic tracking in space (the between island covariance between the intercept and the environment of selection) depends on the PO-regressions and DO-regressions are shown in d) and e) respectively. f) shows how genetic tracking in space depends on spatial parameters when temporal parameters are fixed. The remaining fixed parameter values are *A* = 0, *B* = 1, *G*_*aa*_ = *E*_*aa*_ = 1, *G*_*bb*_ = *E*_*bb*_ = 0, *ω*_*z*_ = 1, *ω*_*b*_ = 3,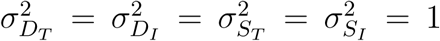 and *S*_*i*_ = 0. From the assumption that environmental fluctuations specific to a time and place are zero, 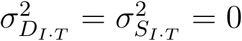. Whenever constant, *κ*_*I*_ = 0.5, *κ*_*T*_ = 0.8, *α*_*I*_ = 0.5 and *α*_*T*_ = 0.5.

In Panel d) of Figure 2, local adaptation increases as the probability of philopatry increases, as expected. Interestingly, for a given level of philopatry, local adaptation also slightly increases as the temporal autocorrelation in the environment of selection increases. This occurs because temporal fluctuations can be tracked genetically, which reduces mean plasticity and therefore promotes local adaptation in space. Panel e) shows that as the spatial DO-regression approaches 0, local adaptation increases, as a consequence of plasticity failing to track spatial fluctuations (because the cue is completely unreliable). Additionally, if the temporal and spatial DO-regressions have the same sign, the plasticity that evolves to cope with temporal fluctuations is also useful, to some degree, for tracking spatial fluctuations and so the amount of local adaptation decreases. When the DO-regressions take their maximum values, plasticity is maximised and local adaptation minimised. In Panel f), temporal parameters are fixed and spatial parameters are allowed to vary. When there is complete philopatry, the PO-regression is 1, local adaptation is maximised and plasticity evolves to cope with temporal fluctuations only. However, the amount of local adaptation still depends to a small degree on the spatial DO-regression. This occurs because the plasticity that evolves in response to temporal fluctuations pushes populations away from their local spatial optima, and this increases as the spatial DO-regression deviates from the temporal DO-regression. Local adaptation is then required to compensate for this and so increases as the spatial DO-regression deviates from the temporal DO-regression.

*Figure 2 here*

### Simulations

In the SI we give a comprehensive assessment of how robust our approximations are, but here we simply choose to show how robust our *G*^*bb*^ *→* 0 approximation for the mean plasticity is across a range of migration rates and strengths of selection on the phenotype, retaining the assumption that 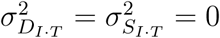. The parameter values that were chosen are the most extreme in terms of breaking our assumptions. In our simulations, we assume *G*^*bb*^ to be equal to *G*^*aa*^, rather than approaching 0.

The accuracy of our approximation is unlikely to be a monotonic function of the width of the fitness function on the phenotype (*ω*_*z*_). When *ω*_*z*_ is small, the strength of selection on the phenotype is strong and so *γ*_*z it*_ is not constant, as we assume, because 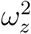 does not dominate 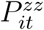 However, this also induces a cost to plasticity in extreme environments because 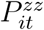 contains the term 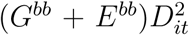 This results in the slope being more constant in time and space and therefore closer to our assumptions. As a consequence, we ran simulations with values of 1, 5, 10 and 20 for *ω*_*z*_. In general, the approximation seems to be accurate, especially when the strength of stabilising selection is weak (*ω*_*z*_ is large) (Figure 3). Standardising *ω*_*z*_ = 20 by the phenotypic variance gives a value of 5.5 which is close to the median value reported in the empirical studies summarised in Kingsolver et al. (2001) (Johnson and Barton, 2005).

**Figure 3:**
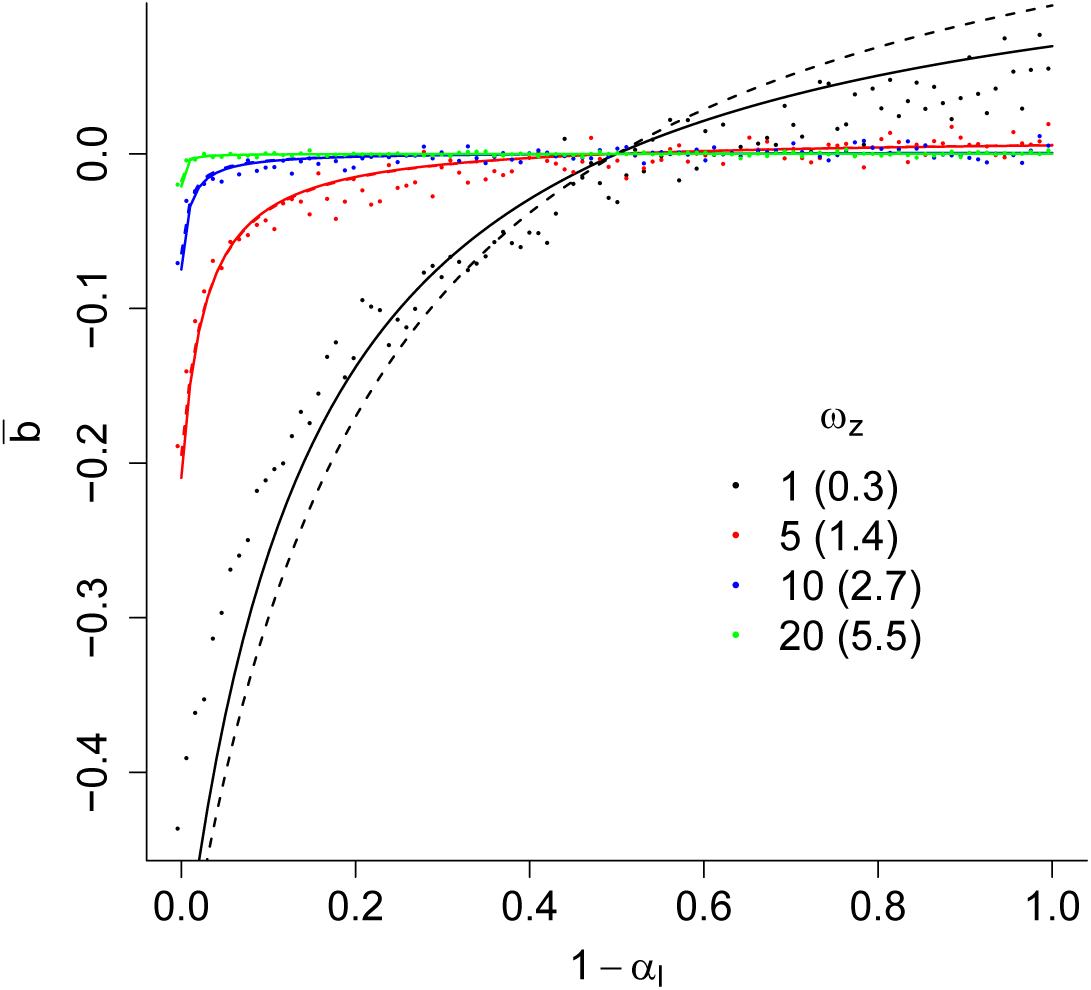
Mean plasticity 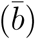 in stochastic simulations with 1,000 islands over 10,000 generations. A single simulation, represented by a single dot, was conducted for each of 100 migration rates (1 *- α*_*I*_), for four different strengths of stabilising selection on the phenotype (*ω*_*z*_; small values indicate stronger stabilising selection), given by the different colours. The number in parentheses is the average value of *ω*_*z*_ scaled by the within-population phenotypic variance. For comparison with the simulations, expected mean plasticities obtained using the approximation *G*^*bb*^ *→* 0, where *γ*_*z*_ is set to 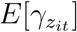, are shown for each strength of stabilising selection and calculated assuming no variance in slopes (dashed line) or a third-order Taylor expansion in *D*_*it*_ (solid line). Parameter values were set to *α*_*T*_ = 0.5, 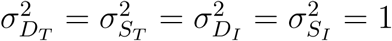, *A* = 0, *B* = 1, *G*^*aa*^ = *E*^*aa*^ = *G*^*bb*^ = *E*^*bb*^ = 1, *κ*_*T*_ = *-*0.8, *κ*_*I*_ = 0.8 and *ω*_*b*_ = 3. From the assumption that environmental fluctuations specific to a time and place are zero, 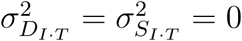.

*Figure 3 here*

### Hyperplasticity and Negative Plasticity

Hyperplasticity in space implies that the regression of the plasticity-induced phenotype 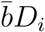 on the environment of selection *S*_*i*_ is steeper than *B*. If *D*_*i*_ and *S*_*i*_ were the same environmental variable this definition reduces to 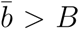 as in (Scheiner and Holt, 2012). Retaining the assumption that fluctuations specific to a time and place are zero, and assuming *B* to be positive, the condition for hyperplasticity to occur is

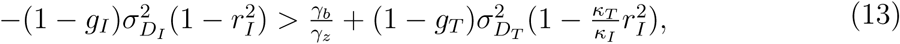

where 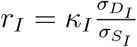 is the spatial correlation between the environments of development and selection and must lie between *-*1 and 1. This equation tells us that spatial hyperplasticity is more likely to occur when genetic tracking is harder in time than in space (*g*_*T*_ *< g*_*I*_), and the regression of the environment of selection on development is steeper in time than in space (*κ*_*T*_ *> κ*_*I*_). However, there must be some association between the environments of development and selection in space 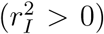 otherwise the plastic response would be flat with respect to spatial variation in the environment of selection. If the environments of development and selection are the same variable but experienced at different times or places then inequality 13 can never be satisfied if autocorrelation is positive.

Negative plasticity implies that the regression of the plastic component of the phenotype on the environment of selection is negative (when *B* is positive). This occurs when

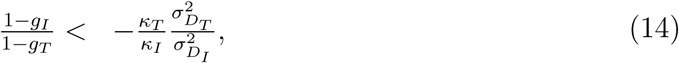

which implies *κ*_*T*_ and *κ*_*I*_ must have different signs, and the temporal association between the environments of selection and development is strong relative to the capacity to adapt in time. Switching the subscripts *I* and *T* gives the equivalent expressions for hyperplasticity and negative plasticity in time (Grether, 2005). In Figure 4, two hypothetical scenarios are illustrated where the conditions for hyperplasticity and negative plasticity are met. In the SI we relax that assumption that fluctuations specific to a time and place do not exist and show the conditions for the evolution of hyper or negative plasticity would be even less stringent since the capacity to adapt to these fluctuations is zero.

**Figure 4:**
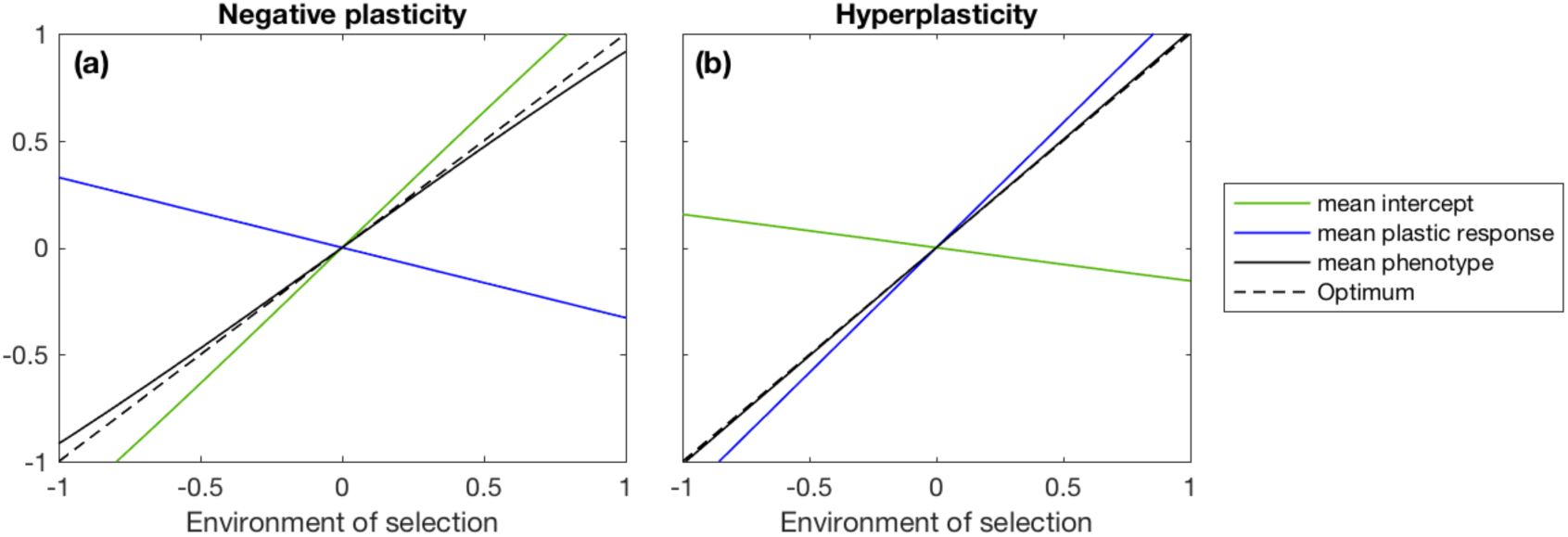
Graphical representations of negative plasticity (a) and hyperplasticity (b) across the average environment of selection of each island. Full lines correspond to island expectations of the intercept (green), effect of plasticity 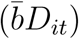 (blue) and phenotype (black), averaged over time, and the dashed line corresponds to the phenotypic optimum. a) shows that when the DO-regression coefficient is negative in time (*κ*_*T*_ = *-*0.8) and positive in space (*κ*_*I*_ = 0.8), plasticity causes a spatial change in phenotype that is opposite in sign to the change in the optimum. Environmental variances are 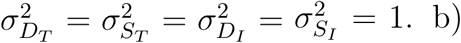 shows that when both the DO-regression coefficient and the environmental variances are greater in time (*κ*_*T*_ = 2 and 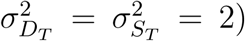) than in space (*κ*_*I*_ = 0.8 and 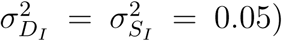), plasticity can evolve to values that overshoot the optimum. In both cases, if the rate of philopatry is high enough (*α*_*I*_ = 0.99), subpopulations can genetically track spatial fluctuations to counteract the effects of plasticity. The remaining fixed parameter values are *A* = 0, *B* = 1, *G*_*aa*_ = *G*_*bb*_ = *E*_*aa*_ = *E*_*bb*_ = 1, *ω*_*z*_ = 1, *ω*_*b*_ = 3, *α*_*I*_ = 0.99 and *α*_*T*_ = 0.5. From the assumption that environmental fluctuations specific to a time and place are zero, 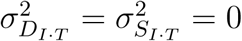

*Figure 4 here*

## Discussion

In this manuscript we show that plasticity evolves in response to spatial variation in the environment in exactly the same way as it does to temporal variation. However, care must be taken to scale any autocorrelation in the environments by dispersal distance and generation-time, respectively. This scaling gives both spatial and temporal autocorrelation the same meaning: the degree to which an individual’s environment is predicted by that of its parents (PO-regression). When this autocorrelation is non-zero, genetic responses to environmental fluctuations are possible and favoured over plasticity when plasticity is costly. When the cost is high and genetic tracking of the environment is easy, the plastic slope tends to zero; but when the cost is low and genetic tracking hard, the slope tends to the regression of the optimum on the environment of development (DO-regression) (Gavrilets and Scheiner, 1993). The plastic slope that evolves is symmetric with respect to temporal and spatial parameters, providing a theoretical basis for the assumptions underlying space-for-time substitutions in empirical work (Wogan and Wang, 2017). However, temporal and spatial fluctuations can have asymmetric influences if there are differences in the values of their homologous parameters, suggesting care must be taken. For example, if the PO-regression is higher and the DO-regression shallower in space than in time, genetic tracking is easier and plasticity less effective in response to spatial than temporal fluctuations. This gives rise to a scenario where the plastic response mainly evolves to cope with temporal fluctuations and tends to the temporal DO-regression in the absence of a cost to plasticity. This can result in spatial hyperplasticity when the evolved plastic slope exceeds the spatial DO-regression, or even in negative plasticity in those rare instances where spatial and temporal DO-regressions have different signs. In these cases, genetic tracking in space acts in the opposite direction to plasticity, resulting in genetic compensation and countergradient variation (Grether, 2005; Levins, 1968; Conover and Schultz, 1995). Whereas previous authors have suggested these patterns are due to maladaptive plasticity, here we show they may be adaptive responses to spatiotemporal variation jointly.

Our conclusions are at odds with previous work looking at the evolution of plasticity when the environment varies in both space and time (Scheiner, 2013). Using simulations, Scheiner (2013) concluded that space and time are not equivalent and plasticity can evolve more easily in response to spatial heterogeneity. The discrepancy between our results and those of Scheiner (2013) arise because in Scheiner (2013) the environments of development and selection are the same variable, often separated by a time lag during which migration may happen. When the environments of development and selection are treated this way, it is very hard to distinguish the effects of migration, generation-time and cue reliability, and important insights can be missed (de Jong, 1999). Our results do recapitulate some of Scheiner’s (2013) findings, such as the evolution of hyperplasticity when both spatial and temporal fluctuations exist. However, here we attribute it to an evolved plastic response to fluctuations in one dimension being maladaptive in the other. This is tolerated when the capacity to genetically track environmental change is stronger in the maladapted dimension because a genetic trend in the opposite direction to plasticity can evolve allowing the phenotype to more closely track the optimum. In contrast, Scheiner (2013) attributed the evolution of hyperplasticity to bet-hedging (Scheiner and Holt, 2012; Scheiner, 2013) although it is unclear whether both spatial and temporal fluctuations would be necessary if the environments of development and selection were allowed to be different. As Tufto (2015) notes, the evolution of bet hedging in these simulations probably arises because there are big fluctuations in the optimum phenotype. This selects for increased phenotypic variance (Bull, 1987) because the population is often in a region where the fitness function is convex and hence disruptive selection predominates (Tufto, 2015). Without a separate mechanism for increasing the phenotypic variance, as in Tufto (2015), a hyperplastic response to an environmental variable can generate this form of bet-hedging (Tufto, 2015; Scheiner, 2013; Scheiner and Holt, 2012). Our approximations ignore this source of selection, although as Tufto (2015) states, the conditions that promote it are probably quite rare in nature.

Genetic trends in space that act in opposition to plastic trends have been called countergradient variation (Conover and Schultz, 1995) and have been demonstrated for several traits in several organisms using reciprocal transplant and common garden experiments (Conover et al., 2009). These experiments are a robust way of assessing whether genetic compensation exists, because phenotypic differences in a common garden should exceed, or be in the opposite direction to, those observed *in situ*. Such experiments do not require the environments of selection and development to be known. Other methods exist that estimate both plastic responses and the environmental sensitivity of selection, although these require identifying the driving environmental variables. These methods have been most widely applied to long-term individual-based data at a single site and are therefore mainly focused on temporal variation. Lay-date in Great tits (*Parus major*) is perhaps the best studied trait in this context and the plasticity-induced phenotype is found to closely track the optimum with no evidence of (temporal) hyperplasticity (Vedder et al., 2013; Gienapp et al., 2013). The optimum in these studies was indirectly estimated using peak caterpillar abundance, but a direct estimate of the environmental sensitivity of selection gave similar results, suggesting that the conclusions are robust (Chevin et al., 2015). An alternative method, using population-level spatiotemporal data, is also able (with caveats) to estimate plasticity and the environmental sensitivity of selection (Phillimore et al., 2010; Hadfield, 2016). Applying this method to Great tit lay-dates, spatial patterns were found to be similar to temporal patterns with little evidence for spatial hyperplasticity (Phillimore et al., 2016). Similar conclusions were drawn using this method for lay-dates of three other passerine birds (Phillimore et al., 2016), but evidence of spatial hyperplasticity for other traits in other taxa is widespread (flowering/leafing time in 4/22 species of plant (Tansey et al., 2017) and most flight dates in 31 species of butterfly (Roy et al., 2015)). However, a drawback of these correlational approaches arises when the driving environmental variables have been misidentified (Michel et al., 2014), or when there are multiple environmental variables but only one has been measured (Chevin and Lande, 2015). It is then possible to obtain spurious estimates that result in the appearance of hyperplasticity or negative plasticity (Chevin and Lande, 2015). Testing whether the hyperplasticity identified in Tansey et al. (2017) and Roy et al. (2015) is real or driven by un/miss-measured variables would require a common garden or reciprocal transplant approach. However, such experiments do not shed light on whether hyperplasticity or negative plasticity is unconditionally maladaptive, as is often believed, or whether it is driven by an adaptive response to temporal fluctuations, as in our model.

Testing whether spatial hyperplasticity is due to the evolution of plasticity to cope with temporal fluctuations probably requires the environments of selection and development to be identified for a trait exhibiting spatial hyperplasticity. If it can be shown that the regression of the environment of selection on development is steeper in time than in space, and spatial autocorrelation in the environment of selection over one dispersal distance is greater than temporal autocorrelation over one generation, then this would be consistent with spatial hyperplasticity being a consequence of adaptive plasticity in response to temporal fluctuations. Alternatively, if individual-based long-term data were available from multiple populations, it would be possible to measure the optimum trait value using fitness and trait data alone. Under this scenario, only the environment of development would need to be identified, and the regression and autocorrelation properties defined above could be framed in terms of optimum trait value (*θ*_*zit*_ = *A* + *BS*_*it*_) instead of the environment of selection (*S*_*it*_). The statistical methodology outlined in Chevin et al. (2015) could be extended to such a situation, but the challenges of obtaining such data would be formidable.

Is it surprising that hyperplasticity is not more commonly observed, given that most aspects of the environment vary both spatially and temporally? The simplest explanation is that the general properties of environmental variation make it unlikely. If the environ-ments of development and selection are the same variable but experienced at different times or places, our analytical results suggest that negative autocorrelation is required, which is probably rare. When the environments of development and selection are different variables we have shown that the conditions for hyperplastcity to evolve are less restrictive, unless the relationship between them is similar in space and time. In this instance, spatial and temporal DO-regressions would be similar, resulting in intermediate plastic slopes in both dimensions. Another possibility is that hyperplasticity is rare because of the properties of organisms, which may be evolved features. In our model, there is only one environment of development and so it is unclear whether evolution would favour the use of other cues if they had different relationships to the environment of selection. It is possible that organisms evolve to respond to spatial and temporal fluctuations in the environment of selection by using several cues that pick up on different aspects of the total variation (Chevin and Lande, 2015). For example, imagine a migratory bird that arrives in the northern hemisphere in mid-April and needs to time its breeding so that some number of degree days have occurred before its chicks hatch in June. Photoperiod in mid-April varies spatially but not interannually, making it a reliable cue for latitudinal differences in spring temperature. However, interannual differences in spring temperatures may be better predicted by temperature on arrival, such that birds use both photoperiod and arrival temperature as a means of extracting independent information about spatial and temporal patterns.

Extending the model to multiple cues would be required in order to get a more comprehensive answer to this question. However, the current model does provide insights into how a single cue that fluctuates in time and space influences the evolution of phenotypic plasticity. Given that most environmental variables that fluctuate in space also fluctuate in time, we hope the model is a more realistic description of how and why plasticity evolves.

## Acknowledgements

We thank Ally Phillimore for extensive discussions, Jarle Tufto & Russ Lande for help with solving some of the equations, and Samuel Scheiner for useful feedback on this work. JDH is supported by a Royal Society Research Fellowship (UF150696). JGK was supported by the University of Edinburgh.

## Author Contributions

JDH designed the study; JGK and JDH conducted the research and wrote the paper.

## Data Accessibility

No new data were created during this study.

